# Comparison of linear combination modeling strategies for GABA-edited MRS at 3T

**DOI:** 10.1101/2021.05.26.445817

**Authors:** Helge J. Zöllner, Sofie Tapper, Steve C. N. Hui, Peter B. Barker, Richard A. E. Edden, Georg Oeltzschner

## Abstract

**Purpose:** J-difference-edited spectroscopy is a valuable approach for the in vivo detection of γ-aminobutyric-acid (GABA) with MRS. A recent expert consensus article recommends linear combination modeling (LCM) of edited MRS but does not give specific details of implementation. This study explores different modeling strategies to adapt LCM for GABA-edited MRS.

**Methods:** 61 medial parietal lobe GABA-edited MEGA-PRESS spectra from a recent 3T multi-site study were modeled using 102 different strategies combining six different approaches to account for co-edited macromolecules, three modeling ranges, three baseline knot spacings, and the use of basis sets with or without homocarnosine. The resulting GABA and GABA+ estimates (quantified relative to total creatine), the residuals at different ranges, SDs and CVs, and Akaike information criteria, were used to evaluate the models’ performance.

**Results:** Significantly different GABA+ and GABA estimates were found when a well-parameterized MM_3co_ basis function was included in the model. The mean GABA estimates were significantly lower when modeling MM, while the CVs were similar. A sparser spline knot spacing led to lower variation in the GABA and GABA+ estimates, and a narrower modeling range – only including the signals of interest – did not substantially improve or degrade modeling performance. Additionally, results suggest that LCM can separate GABA and the underlying co-edited MM_3co_. Incorporating homocarnosine into the modeling did not significantly improve variance in GABA+ estimates.

**Conclusion:** GABA-edited MRS is most appropriately quantified by LCM with a well-parameterized co-edited MM_3co_ basis function with a constraint to the non-overlapped MM_0.93_, in combination with a sparse spline knot spacing (0.55 ppm) and a modeling range between 0.5 and 4 ppm.

**Graphical Abstract:** 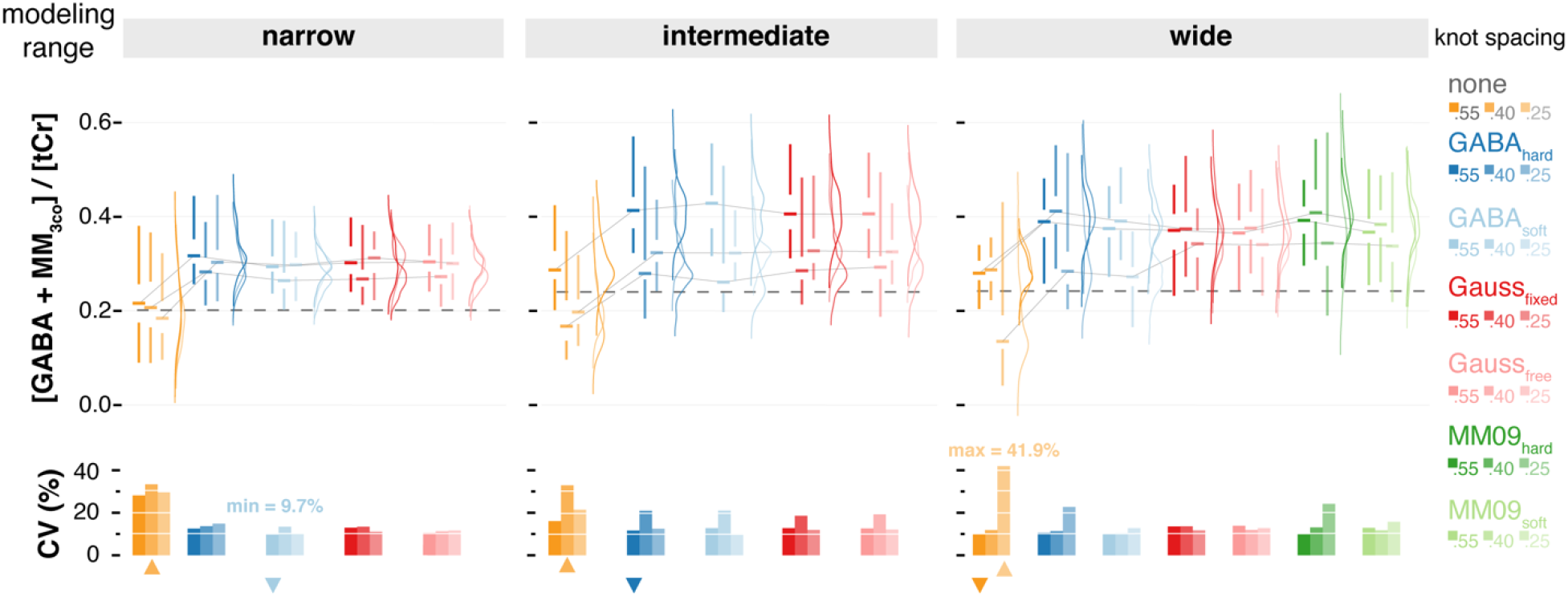

102 strategies to model GABA-edited MRS with linear combination modeling were evaluated to quantify GABA and GABA+ in Osprey. Significantly different GABA and GABA+ estimates were found when a well-parameterized macro-molecule at 3 ppm was included. The findings suggest that linear combination modeling needs to be adapted for quantification of GABA-edited MRS.

## Introduction

A recent expert consensus paper recommended that linear combination modeling (LCM) should be used for the quantification of edited MRS data^1^, stating that standard fitting approaches originally optimized for short-TE MRS should be adapted for edited MRS. Further, it was recommended that quantum-mechanical simulations should be used to confirm the co-edited profile of all metabolites in the edited spectrum, and contributions from macromolecule (MM) signals should be specified. Despite these recommendations, little detail was given regarding several unique features of edited spectra, and how they should be appropriately modeled. These features include:

1. The MEGA-PRESS experiment is well-known to co-edit MM signals with coupled spins at 1.7 and 3 ppm, causing substantial contamination of the edited GABA signal, and forcing researchers to report the composite measure GABA+MM (GABA+)^1^. Because the co-edited MM signal is poorly characterized, there is currently no consensus or recommendation on how to appropriately account for it during spectral modeling. Instead, the most widely used analysis algorithms implement entirely different strategies to fit the composite 3-ppm signal. For example, the Gannet software uses a single Gaussian model^2^, while a double-Gaussian is used in Tarquin^3^, and LCModel^4^ defaults to a basis set that only includes the GABA basis function.
2. Another co-edited compound contributing to the 3 ppm signal is homocarnosine (HCar), a dipeptide of GABA and histidine. While the 3 ppm multiplets of GABA and homocarnosine are separated by just 0.05 ppm (which are therefore unlikely to be successfully separated), inclusion of a homocarnosine basis function may be warranted based on its reported concentration in vivo (∼0.5 mmol/kg ^5^, compared to ∼1-2 mmol/kg for GABA), but it has not been investigated whether doing so has a stabilizing or destabilizing effect on the modeling^6^.
3. Unedited spectra are typically modeled over a restricted frequency-domain range covering the visible upfield peaks, including macromolecular and lipid resonances between 0 and 1 ppm, but usually avoiding the water suppression window above ∼4 ppm. The choice of frequency-domain modeling range for edited spectra is less obvious. Since the main advantage of spectral editing is the isolation of a single target resonance, modeling signals outside the immediate surrounding of the target may dilute the resolving power of editing. On the other hand, increasing the modeling range may offer useful constraints to stabilize the solution of the modeling problem. The difference is highlighted by the different strategies encountered in common software tools – while the Gannet software fits the GABA-edited difference spectrum over a narrow range (only including the 3-ppm GABA+ and 3.75 ppm glutamate and glutamine peaks), the LCModel recommendation is to include the strong co-edited signals from glutamate (Glu), glutamine (Gln), glutathione (GSH), N-acetylaspartylglutamate (NAAG), and N-acetylaspartate (NAA), which heavily overlap with GABA around 2.25 ppm. The effects of limiting the modeling range have not been assessed systematically to date.
4. Linear combination modeling methods commonly include terms to account for smooth baseline curvature, usually parametrized from cubic B-spline or polynomial functions, or by smoothing residuals. The flexibility of the baseline model substantially affects metabolite estimates from unedited spectra^7^; while baseline terms are necessary to account for e.g. lipid contamination, poor water suppression etc., they are potential sources of overfitting if awarded too many degrees of freedom. Baseline modeling may have an even greater influence when modeling difference spectra, since only *co-edited* lipid and MM signals contribute to the smooth background variation. Importantly, the co-edited MM background of the GABA-edited difference spectrum has not been appropriately characterized (e.g., through metabolite-nulled acquisition), suggesting that the choice of baseline flexibility can drastically influence modeling results through two highly susceptible regions of the spectrum. First, in the absence of an appropriate model for the co-edited broad MM signal at 3 ppm, this signal may be absorbed into the baseline depending on its flexibility. Second, strong MM and lipid signals in the region between 0.5 and 2.5 ppm may be affected by the 1.9 ppm editing pulse (either directly through saturation or indirectly through coupling), likely leading to an unknown, but substantial, MM contribution in this spectral region^8,9^. This is especially important considering that the co-edited signals from NAA, NAAG, Glu, Gln, and GSH overlap with GABA in this region. Overly rigid baselines may provide insufficient flexibility to capture these signals, in turn compromising the accuracy of the estimation of the co-edited metabolites.

The aim of this study was to evaluate different strategies for linear combination modeling of GABA-edited MEGA-PRESS difference spectra, and to establish initial ‘best practices’ substantiating the recommendations of the expert consensus on spectral editing. To this end, different approaches to account for co-edited MM signals, various modeling ranges and baseline knot spacings, as well as the inclusion of homocarnosine were compared. In the absence of a ‘gold standard’, the performance of each modeling strategy was assessed by comparing descriptive statistics of the metabolite estimates, calculating the Akaike information criteria, and assessing the fit residuals.

## Methods

### Study participants & data acquisition

In this study, 61 publicly available GABA-edited MEGA-PRESS datasets originating from 7 sites from a recent 3T multi-center study^10^ were analyzed (see **Supplementary Material 1** for subject list). All datasets were acquired on Philips 3T scanners with the following acquisition parameters: TR/TE = 2000/68 ms; 320 excitations (10m 40s scan time); 16-step phase-cycle; 2 kHz spectral width; 2000 samples; 27-ml cubic voxel volume in the medial parietal lobe. For this heuristic approach of exploring the GABA modeling, the data homogeneity (SNR, FWHM, tissue composition, and absence of fat contamination) was increased while reducing the overall number of subjects by including only 61/298 subjects of the original dataset^10^. All sites except for P8 used a similar sequence implementation with interleaved water referencing for prospective frequency correction^11^. For the edit-ON transients, the editing pulses with 15 ms pulse duration and 82.5 Hz inversion bandwidth (FHWM) were applied at a frequency of 1.9 ppm to refocus the coupling evolution of the GABA spin system. For the edit-OFF transients, the editing pulses were applied at a frequency of 7.5 ppm. Edit-ON and edit-OFF transients were acquired in alternating order. An additional water reference scan was acquired for each dataset using interleaved water referencing ^11^, i.e. one excitation with water suppression and editing pulses deactivated every 40 water-suppressed excitations (total of 8 averages).

### Data pre-processing

Data were analyzed in MATLAB using Osprey^12,13^ (v.1.0.1.1), a recently published open-source MRS analysis toolbox. Raw data were eddy-current-corrected ^14^ based on the water reference, and individual transients were aligned separately within the edit-ON and edit-OFF conditions using the robust spectral registration algorithm^15^. Averaged edit-ON and edit-OFF spectra were aligned by optimizing relative frequency and phase such that the water signal in the difference spectrum was minimized. The final difference spectra for quantification were generated by subtracting the edit-OFF from the edit-ON spectra. Finally, any residual water signal was removed with a Hankel singular value decomposition (HSVD) filter^16^ to improve data quality in the edit-OFF spectra and to reduce residual baseline roll in the difference spectra.

### Basis set

The basis set used for modeling was generated from a fully localized 2D density-matrix simulation of a 101 x 101 spatial grid (voxel size: 30 mm x 30 mm x 30 mm; field of view: 45 mm x 45 mm x 45 mm) implemented in a MATLAB based simulation toolbox FID-A ^17^, using vendor-specific refocusing pulse shape and duration, sequence timings, and phase cycling. It contains 17 metabolite basis functions (ascorbate, aspartate, creatine (Cr), negative creatine methylene (- CrCH_2_), GABA, glycerophosphocholine, GSH, Gln, Glu, water, myo-inositol, lactate, NAA, NAAG, phosphocholine, phosphocreatine (PCr), phosphoethanolamine, scyllo-inositol, and taurine) and 8 Gaussian MM and lipid resonances (MM_0.94_, MM_1.22_, MM_1.43_, MM_1.70_, MM_2.05_, Lip09, Lip13, Lip20, details in **Supplementary Material 2** with similarly defined parametrization as described in the LCModel software manual^18^) for the edit-OFF spectrum.

For the difference spectrum, MM_0.94_ and the co-edited macromolecular signal at 3 ppm (MM_3co_) were parametrized as Gaussian basis functions (MM_0.94_: 3-proton signal; chemical shift 0.915 ppm, full-width at half-maximum (FWHM) 11 Hz; MM_3co_: 2-proton signal; chemical shift 3 ppm; FWHM 14 Hz). The MM_3co_ amplitude was defined under the assumptions of a pseudo-doublet GABA signal at 3 ppm and the MM_3co_ contribution to the 3-ppm GABA peak to be around 50%^1,6,8,19^. The optimum FWHM used to parametrize the MM_3co_ basis function was determined to be 14 Hz by fitting the mean difference spectrum of all datasets with a composite GABA+ basis function (GABA + MM_3co_) with varying FHWM (between 1 and 20 Hz). The parameterized Gaussian MM_3co_ basis function was integrated into the modeling process using different assumptions and constraints described in the following paragraphs.

### Linear combination modeling of GABA-edited difference spectra

Osprey’s frequency-domain linear combination model was used to determine the metabolite estimates. Model parameters include metabolite basis function amplitudes, frequency shifts, zero/first order phase correction, Gaussian and Lorentzian linebroadening, and cubic spline baseline coefficients. All parameters are determined by Levenberg-Marquardt^20,21^ non-linear least-squares optimization, using a non-negative least-squares (NNLS) fit ^22–24^ to determine the metabolite amplitudes and baseline coefficients at each iteration of the non-linear optimization. Amplitude ratio soft constraints are imposed on MM and lipid amplitudes, as well as selected pairs of metabolite amplitudes, as defined in the LCModel manual^4,18^. The strength factor of the amplitude ratio soft constraint λ is set to 0.05 by default.

A range of modeling strategies for the GABA-edited difference spectrum was included in this study, covering various aspects of the modeling process (**Figure 1**). The different parametrizations and soft constraints to account for the co-edited MM_3co_ signal are shown in **Figure 1A**. All possible combinations for the modeling strategies: i) inclusion of homocarnosine in the basis set; ii) different modeling ranges; iii) different baseline spline knot spacings and iv) different parametrizations and soft constraints to account for the co-edited MM_3co_ signal are tabulated graphically in **Figure 1 B**. Each modeling aspect is described in detail below:

**Figure 1.**
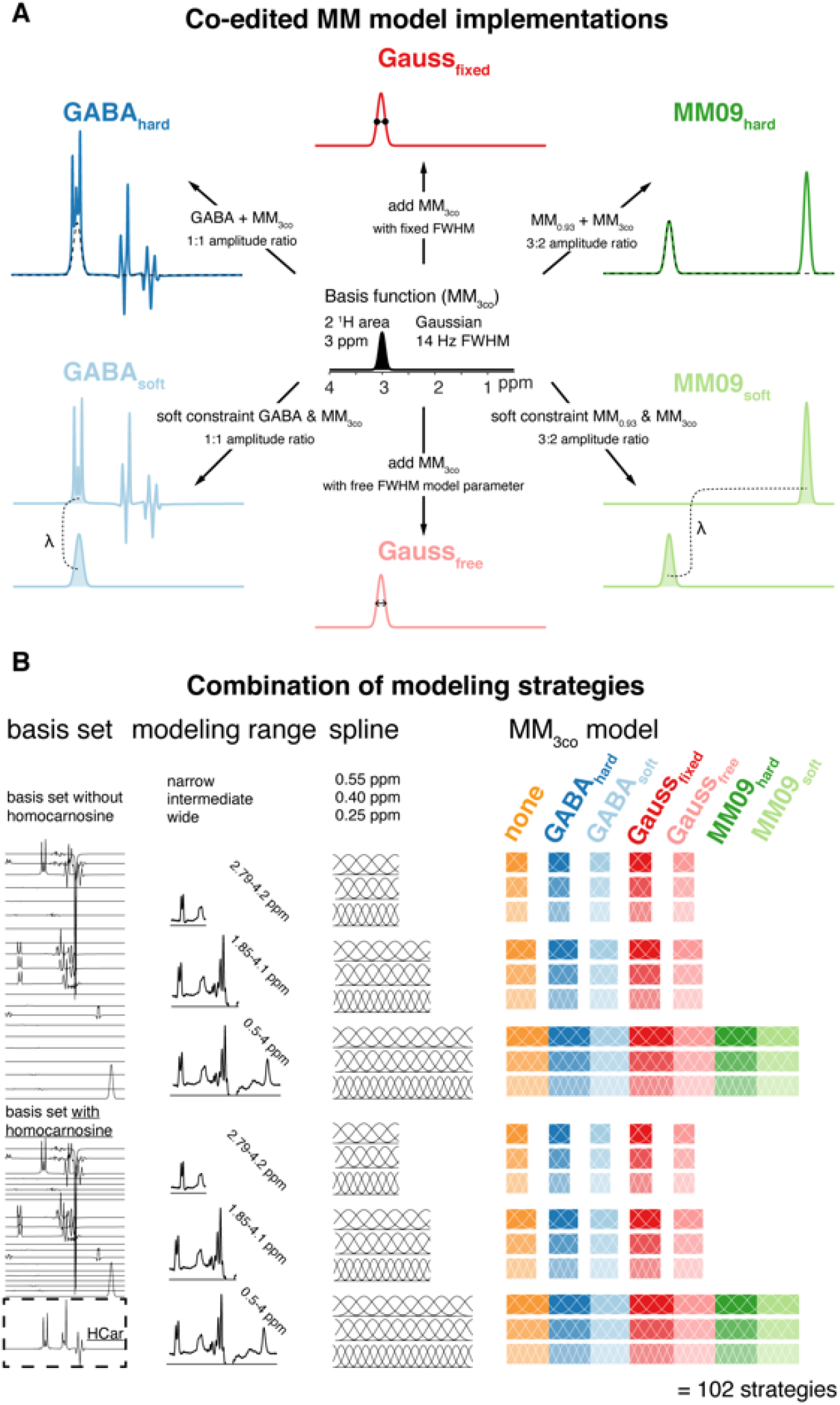
Different linear combination modeling strategies for GABA-edited spectra. (A) Different co-edited MM_3co_ modeling approaches derived from a Gaussian function at 3.0 ppm (B) All combinations of basis set composition, modeling range, spline knot spacing, and MM_3co_ modeling leading to 102 different modeling strategies.

### Including homocarnosine in the basis set

To assess the effects of including homocarnosine in the linear combination model, we repeated all analysis steps with two different basis sets: the default Osprey basis set *with* and *without* an additional HCar basis function. Chemical shift and scalar coupling parameters describing the HCar spin system were taken from literature^6^.

### Varying the modeling range and baseline knot spacing

Two aspects of linear combination modeling are suggested to have a considerable influence on metabolite estimates^7,25^. First, the choice of the modeling range, i.e., the frequency interval that defines the part of the frequency-domain spectrum that is considered to calculate the least-squares difference between model and data. Second, the baseline knot spacing, i.e., the frequency difference between two adjacent knots of the cubic spline basis that is used to approximate the smooth baseline.

Three different modeling range scenarios were considered, reflecting common choices in the literature and widely used software tools: a) a wide modeling range typically used to analyze unedited spectra, including all signals in the GABA-edited difference spectrum (0.5 to 4 ppm – *“wide”*); b) an intermediate modeling range excluding signals below 1.9 ppm (e.g. co-edited lipids and macromolecules), but including strong co-edited signals from NAA, NAAG, Glu, Gln, and GSH (1.85 and 4.1 ppm, *“intermediate”*), similar to the range recommended in LCModel’s dedicated ‘mega-press-3’ option; and c) a narrow modeling range only including the co-edited signals from GABA+ and Glx (2.79 – 4.2 ppm, *”narrow”*), the default modeling range in Gannet^2^.

Three spline knot spacings were included in the analysis, with 0.4 ppm being the default Osprey option, shown to create reproducible and comparable metabolite estimates for conventional MRS ^26^, as well as sparser (0.55 ppm) and denser (0.25 ppm) spline knot spacings.

### Co-edited macromolecule models

Seven different strategies to model the GABA-edited difference spectrum were implemented (**Figure 1 A**). The trivial approach – not accounting for the co-edited signal MM_3co_ at all – is labeled **none**. The other six modeling strategies all include a dedicated parametrized Gaussian MM_3co_ basis function. This basis function is given different degrees of freedom in the different strategies, e.g., hard- or soft-constrained relative to the amplitude of the GABA or the MM_0.94_ basis functions, and with a fixed or free width. Here, strategies with fewer degrees of freedom reflect the frequently made assumption that the GABA-to-MM ratio (and the MM background itself) is relatively stable across subjects and anatomical region, and assumed to be known, while strategies with more degrees of freedom or soft constraints relax these assumptions:

- The **GABA_hard_** model uses a single composite GABA+MM basis function by adding the GABA and MM_3co_ (initial FWHM of the basis function = 14 Hz) basis functions with a fixed 1:1 amplitude ratio. The 1:1 ratio reflects the widely used empirical assumption that 50% of the 3-ppm signal in a conventional GABA-edited difference spectrum can be attributed to co-edited macromolecules^6,19^.
- The **GABA_soft_** model uses separate GABA and MM_3co_ (initial FWHM of the basis function = 14 Hz) basis functions and imposes a soft constraint on the ration of the amplitudes of both basis functions during the optimization (1:1 ratio).
- The **Gauss_fixed_** model uses separate GABA and MM_3co_ (initial FWHM of the basis function = 14 Hz) basis functions. No further constraints are imposed. This means possible changes in the contributions to the 3-ppm GABA peak are modeled.
- The **Gauss_free_** model uses separate GABA and MM_3co_ basis functions. In contrast to the Gauss_fixed_ model, the FWHM of the Gaussian MM_3co_ signal is represented by an additional model parameter. This means that the MM_3co_ basis function itself is not static, but dynamically modified during optimization.
- The **MM09_hard_** model uses separate GABA and MM basis functions. The MM_3co_ basis function is replaced by a composite MM_0.94_ + MM_3co_ basis function (i.e., the MM_0.94_ (initial FWHM of the basis function = 11 Hz) and MM_3co_ (initial FWHM of the basis function = 14 Hz) basis functions are added in a 3:2 ratio). The result is a single composite basis function for MM_0.94_ and MM_3co_, adapted from the soft constraint model described in the literature ^9^.
- The **MM09_soft_** model uses separate GABA, MM_0.94_ and MM_3co_ basis functions. In contrast to the MM09_hard_ model, soft constraints enforce a ∼3:2 amplitude ratio for the MM_0.94_ and MM_3co_ amplitudes during optimization. The use of two separate but linked basis functions for MM_0.94_ and MM_3co_ is similar to previously described implementations ^9^.

The models MM09_hard_ and MM09_soft_ ^27^ as well as Gauss_fixed_ ^28^ correspond to models previously investigated using the LCModel software and the amplitude assumptions were derived empirically. It is worth repeating here, that each basis function receives a separate Lorentzian linebroadening, frequency shift, and amplitude parameter during the optimization, in addition to the global parameters (zero/first order phase correction, global frequency shift, and Gaussian linebroadening). For the Gauss_free_ model, the MM_3co_ basis function is dynamically updated as an explicit modeling parameter during the optimization, therefore the MM_3co_ basis function has effectively two separate adjustable parameters to account for its linewidth (the Lorentzian linebroadening term and the FWHM of the MM_3co_ basis function). Finally, the composite models GABA_hard_ that lacks separate GABA and MM functions has only one linebroadening, one frequency, and one amplitude parameter compared to twice the parameters for its soft constraint counterparts.

Combining the various MM_3co_ models (5 + 2 that were used for the wide modeling range only), modeling ranges (3), baseline spline knot spacings (3), and basis sets (2), a total of 102 different modeling strategies were investigated in this study. All models were implemented in Osprey^12^ and are available on GitHub^13^.

### Quantification, visualization, and statistics

#### Quantification

For the basis set without homocarnosine, GABA refers to the model amplitude estimate for the GABA basis function, which is of course only available for the modeling strategies with separate basis functions for GABA and MM_3co_ (none, GABA_soft_, Gauss_fixed_, Gauss_free_, MM09_soft_). GABA+ refers to the sum of the amplitude estimates for GABA and MM_3co_ (GABA_soft_, Gauss_fixed_, Gauss_free_, MM09_hard_, MM09_soft_) or the amplitude estimate for the composite basis function including both MM and GABA (GABA_hard_) and is therefore calculated for all strategies with an explicit MM_3co_ model. For comparison, the GABA amplitude for the ‘non’ strategy is included in the figures reporting GABA + MM_3co_. However, it still refers to a GABA-only estimate. For the basis set that included homocarnosine (HCar), the difference in GABA and MM_3co_ estimates between the modeling strategies with and without HCar (⊗GABA and ⊗MM_3co_, respectively) were investigated to evaluate whether the inclusion of HCar has a systematic effect on the estimation of those signals with which it overlaps. All estimates were quantified relative to the total creatine (Cr + PCr) amplitude from the edit-OFF spectrum with the wide modeling range and a spline knot spacing of 0.4 ppm. Differences in GABA(+)/tCr between modeling strategies are therefore only related to the modeling of the difference spectra, but not to the reference compound modeling. No further tissue or relaxation corrections were applied. Further, the relative contributions of MM_3co_ to the GABA+ estimate and the relative contributions of HCar to the sum of GABA+ and HCar estimate were calculated.

#### Visualization

The modeling performance and systematic characteristics of each modeling strategy were visually assessed through the mean spectra, mean fit, mean residual, and mean models of GABA+, GABA, MM_3co,_ HCar (if included) and the baseline, i.e., averaged across all datasets.

The metabolite estimate distributions were visualized as violin plots including boxplots with median, 25^th^/75^th^ quartile ranges, and smoothed distributions to identify systematic differences between modeling strategies. In addition, the mean value of the ‘none’ model across the three spline knot spacings was added for each modeling range as a dashed horizontal line. Bar plots were created to visualize quality metrics, including the standard deviation if appropriate. All plots were generated with R^29^ (Version 3.6.1) in RStudio (Version 1.2.5019, RStudio Inc.) using SpecVis^26,30^, an open-source package to visualize linear combination modeling results with the ggplot2 package^31^. All scripts and results are publicly available^32^.

#### Statistics

Significant differences in the mean and the variance of the GABA, GABA+, and MM_3co_ estimates were assessed between all modeling strategies. The statistical tests were set up as paired without any further inference. Differences of variances were tested with Fligner-Killeen’s test, with a post-hoc pair-wise Bonferroni-corrected Fligner-Killeen’s test. The means were compared with an ANOVA or a Welch’s ANOVA, depending on whether variances were different or not. Post-hoc analysis was performed with a paired t-test with equal or non-equal variances, respectively.

Additionally, Pearson’s correlation was used to investigate the impact of including HCar in the basis set. The strength of the correlation was considered substantial for R > 0.25.

#### Model evaluation criteria

The performance of each modeling strategy was evaluated in different ways, including the impact of the different modeling strategies on the GABA, GABA+, and MM_3co_ estimates, as well as several quality measures:

1. Visual inspection: Mean model, residual, and baseline were assessed for characteristic features.
2. SD fit quality: The SD of the residual was determined, and then normalized by the noise level (calculated as the SD of the noise between -2 and 0 ppm). This was done over the entire modeling range of the difference spectrum and termed **residual_SD range_**.
3. Amplitude fit quality: the difference between the maximum and minimum of the residual was determined, and then normalized by the noise level ^25^ (similarly calculated as in the second criterion).This was done over the entire modeling range of the difference spectrum and termed **residual_ampl range_**.
4. Amplitude 3-ppm peak fit quality: Similar to the third criterion, the residual was calculated over the range of 3.027 ± 0.15 ppm to assess the fit quality of the 3-ppm GABA peak and termed **residual_ampl 3ppm_**.
5. Consistency of metabolite estimates: The across-subject coefficients of variation (CV = SD/mean) for all metabolite estimates (GABA/tCr, GABA+/tCr) were calculated for each modeling strategy.
6. Akaike Information Criterion (AIC): The Akaike information criterion ^33^, which takes the number of model parameters into account, is defined as follows:

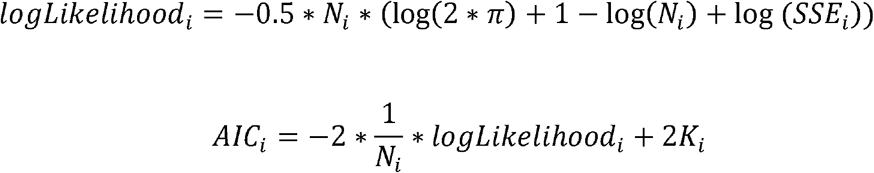

Here, *N_i_* is the number of points in the modeling strategy *i*, *SSE_i_* is the sum of squared error (i.e., squared residual) of that strategy, and *K_i_* is the number of free model parameters for that strategy. The *logLikelihood_i_* was divided by the number of points *N_i_*_t_ to reduce the strong weighting of the datapoints and to make the *AIC_i_* values comparable for different modeling ranges. Soft constraint model parameters were included with a value of 0.5. Lower *AIC_i_* values indicate a more appropriate model. Subsequently, **Δ*AIC_i_*** scores were calculated as the difference of *AIC_i_* of modeling strategy *i* and the model with the lowest *AIC_min_*:

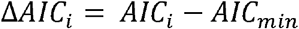

## Results

All 61 datasets were successfully processed and modeled with all 102 modeling strategies. No data were excluded from further analysis. The data quality assessment indicated consistently high spectral quality for all spectra (NAA_SNR_ = 272 ± 70; NAA_FWHM_ = 5.29 ± 1.09 Hz) without lipid contamination. Individual spectra as well as the mean spectrum and SD are displayed in **Supplementary Material 3**. Other data quality measures were extracted from the Gannet^2^ analysis performed in recent multi-site studies^10,34^. For the Philips-only subset of datasets in the present study, the tissue composition (fGM = 0.60 ± 0.04; fWM = 0.27 ± 0.03; fCSF = 0.13 ± 0.04) and across-subject CV (GABA+/Cr = 9.99%) indicate consistency in the dataset and the modeling. Across-subject CV was interpreted as a measure of modeling performance, assuming that increased CVs are mainly introduced by variability in the modeling and do not reflect biologically meaningful variance of GABA+ estimates.

### Summary and visual inspection of the modeling results

**Figure 2** shows the mean modeling results for all modeling strategies without homocarnosine. Not including MM_3co_ leads to a substantial structured residual around 3 ppm for all knot spacings and modeling ranges. In contrast, all modeling strategies with MM_3co_ appear to reflect the lineshape of the 3-ppm signal more accurately, with very similar results for the complete fit (metabolites, MMs, and baseline) and the individual components. Modeling strategies with the intermediate and wide modeling range further show strong residuals around 2 ppm, suggesting slightly inaccurate lineshape modeling of the methyl singlets from NAA and NAAG, or inaccurate modeling of co-edited MM signals in this region. Structured residuals appear also in the region of the 3.75 ppm Glx signals, although they are much less pronounced in strategies with the narrow modeling range, suggesting that including the 2.25 ppm multiplets (and underlying baseline fluctuation) has a considerable impact on phase estimation.

**Figure 2.**
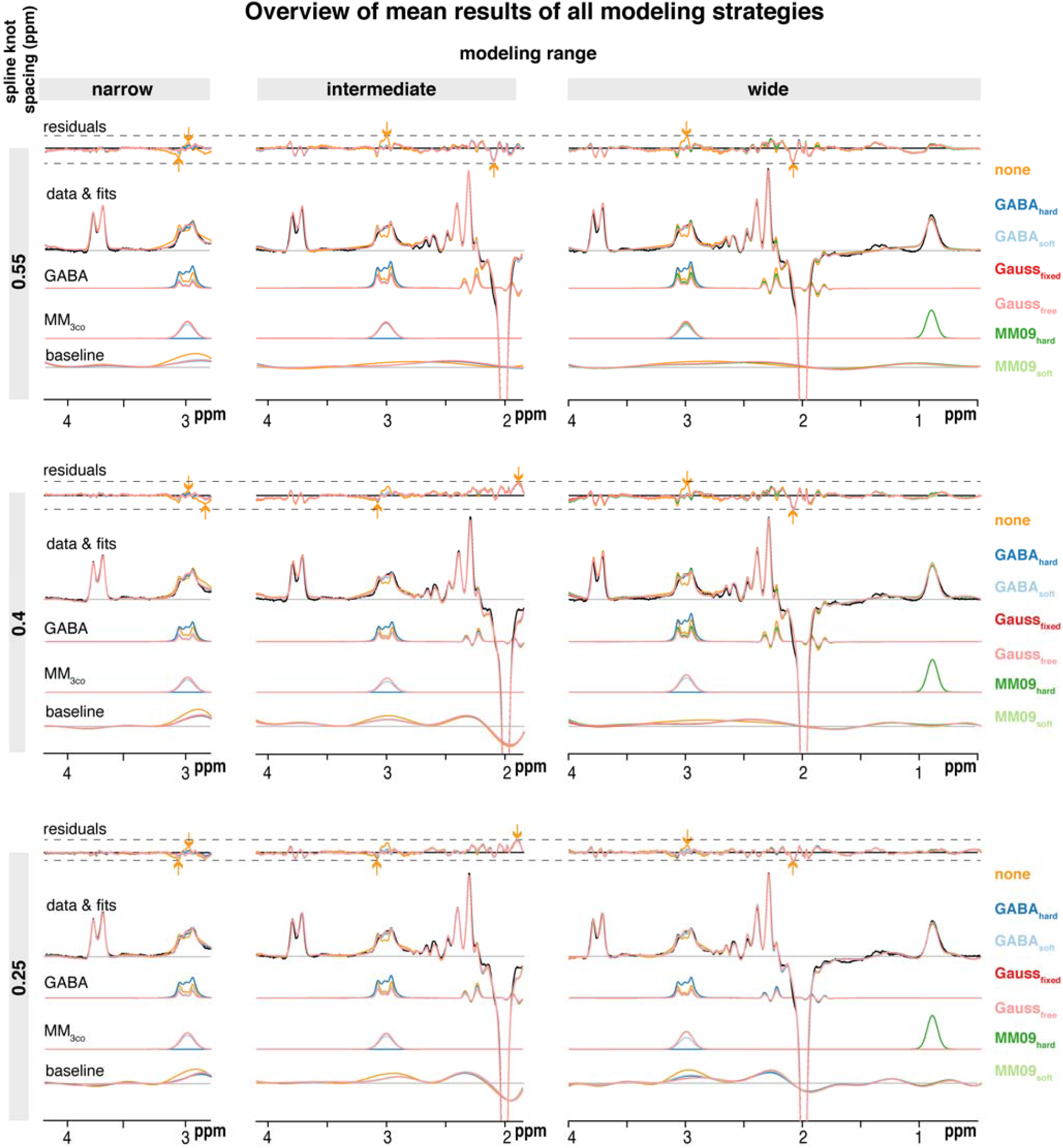
Mean modeling results for all modeling strategies without homocarnosine. A substantial structured residual is apparent at 3 ppm if no MM modeling strategy is included. All three modeling ranges (columns), three spline knot spacings (rows), and MM_3co_ model (color-coded) are presented with mean residuals and fits, as well as the GABA+, GABA, MM_3co_, and spline baseline models. The mean data is included in black. The dashed lines indicate the range of the residual across one row. The arrows indicate the range of values for a specific modeling range and spline knot spacing with the color corresponding to the MM_3co_ model with minimum/maximum value.

In general, the residuals are consistent between different MM_3co_ models for any given knot spacing and modeling range. Notably, residuals tend to be smaller on an absolute scale for denser knot spacing and narrower modeling range.

Mean GABA models agree well between all strategies with a separate MM_3co_ model. The GABA_hard_ strategy appears to produce a larger signal as its GABA basis function includes the MM_3co_ signal, but does not model it separately, while the strategies that do so produce comparable mean MM_3co_ models.

The mean baseline is consistently flatter around 3 ppm for modeling strategies with an explicit MM_3co_ model, while absorbing substantially more signal for the ‘none’ approach without an MM model. This behavior is particularly obvious for the dense knot spacing (0.25 ppm) over the wide modeling range. Baseline curvature generally increases for denser knot spacings around 2.2 ppm for the intermediate and wide range.

### Metabolite level distribution

**Figure 3** shows distributions and coefficients of variation (CVs) of the GABA+ estimates for all modeling strategies. Table 1 summarizes the mean and SD GABA/GABA+ estimates as well as the statistics. GABA+ estimates are significantly higher than GABA-only estimates of the ‘none’ modeling strategy for all modeling ranges and knot spacings, supporting the notion from **Figure 2** that not including an MM model leaves a considerable fraction of the edited 3-ppm signal unmodeled, resulting in substantial residuals or increased baseline amplitude flexion.

**Figure 3.**
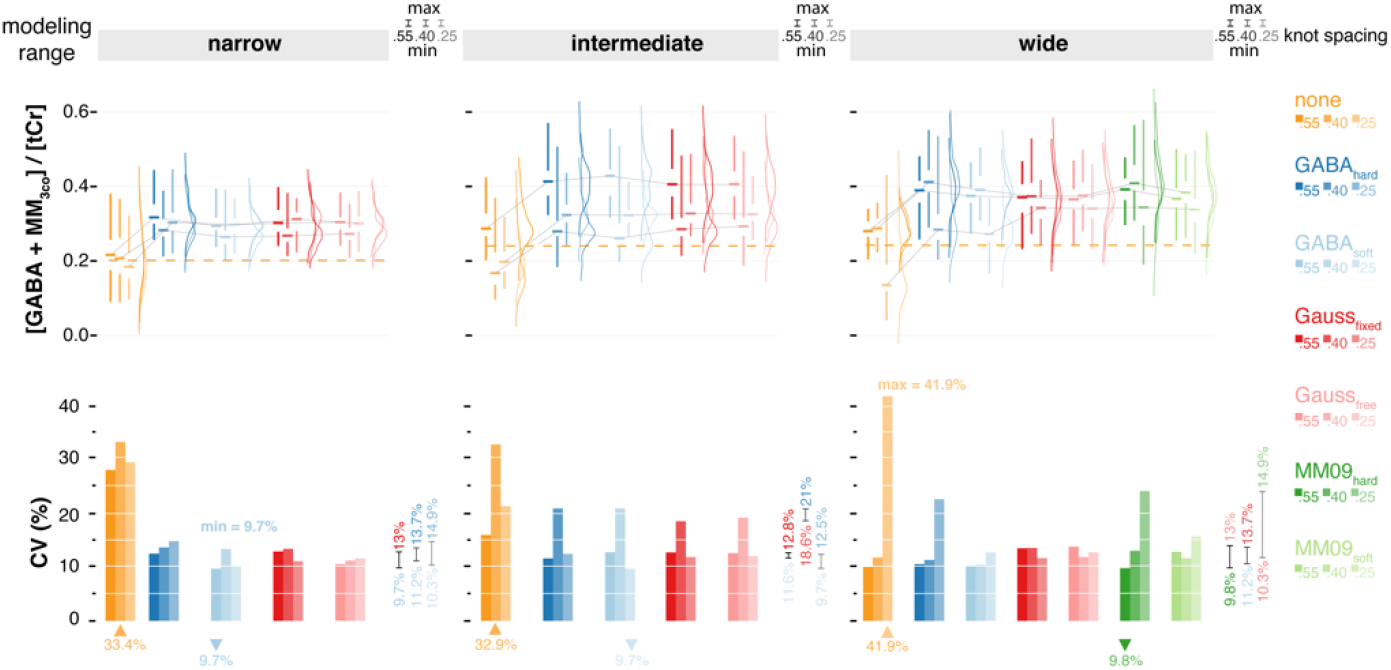
Distribution and across subject coefficients of variation (CVs) of GABA+ estimates for all modeling strategies. Including a MM_3co_ model significantly increases the mean estimates for all modeling strategies, while giving similar or reduced CVs. The mean estimates across the three spline knot spacings of the ‘none’ approach are indicated as a dashed line for each modeling range. All three modeling ranges (column) and three spline knot spacings (within each column), and co-edited MM models (color-coded) are presented. Distributions are shown as half-violins (smoothed distribution), box plots with median, interquartile range, and 25^th^/75^th^ quartile. The median lines of the box plots are connected to visualize trends within a specific baseline knot spacing. CVs are summarized as bar plots. Minimum/maximum CVs for each modeling range are indicated as downwards/upwards triangles in the color corresponding to the MM_3co_ model. Minimum/maximum CVs for each baseline knot spacing within a specific modeling range are reported on the right side of each column. Global minimum and maximum CVs across all models are added as text.

Notably, all modeling strategies with MM_3co_ return comparable mean estimates and CVs within the same knot spacing (see Minimum/Maximum column of **Figure 3**). In addition, sparser knot spacing leads to lower CVs. The intermediate modeling range does not appear to perform more consistently than both other modeling ranges.

### Model evaluation

**Figure 4** summarizes the metrics used for model evaluation. The residual over the modeled frequency range (residual_SD range_ and residual_ampl range_) is lowest for the narrow modeling range. For the intermediate and wide modeling ranges, residual_ampl range_ is substantially higher, largely driven by the 2-ppm region (see also **Figure 2**). Consequentially, residual_ampl range_ is comparable between MM modeling strategies for a given knot spacing (see Minimum/Maximum column of **Figure 4**).

**Figure 4.**
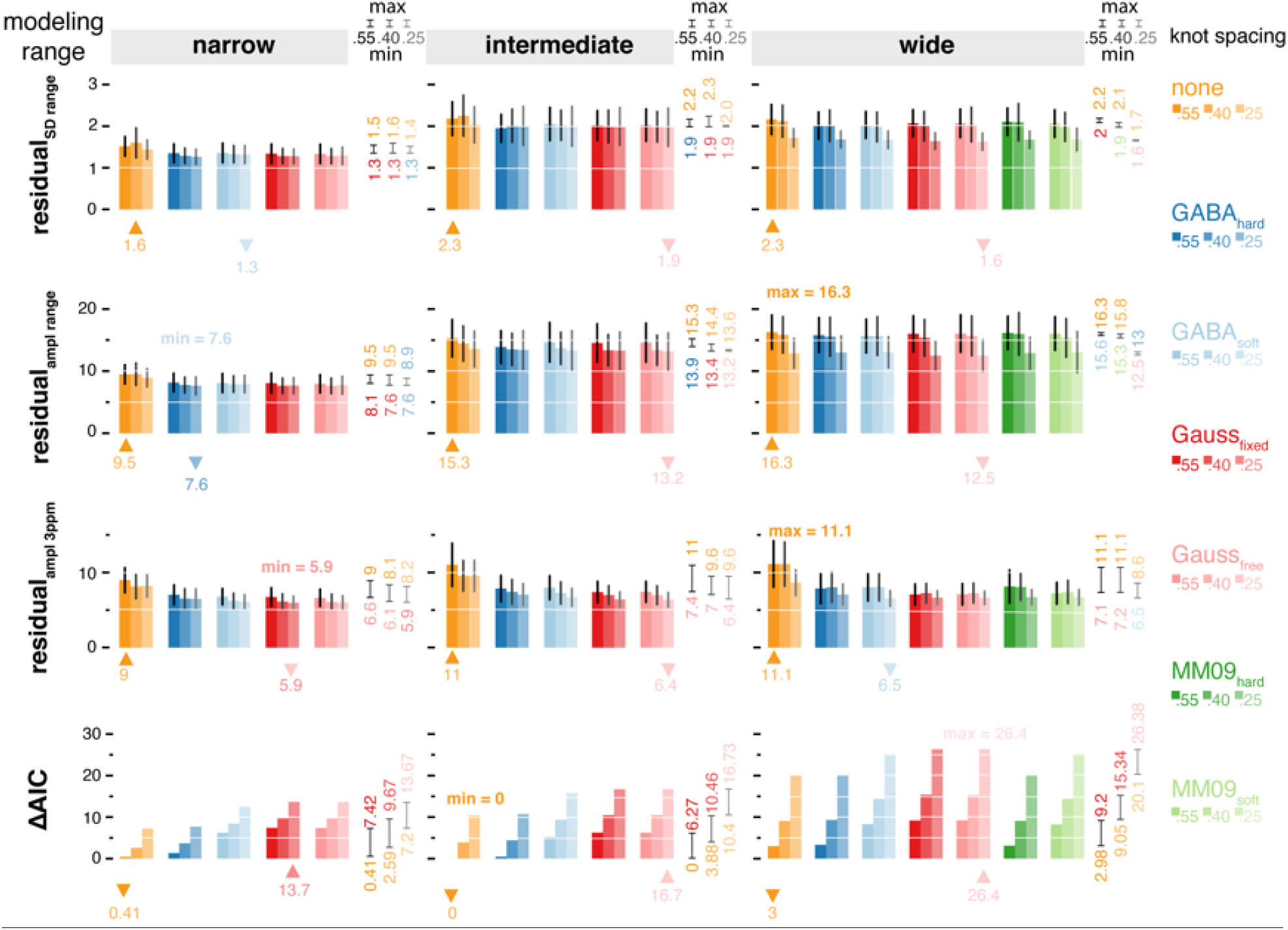
Evaluation of all modeling strategies. Comparably low residual_ampl range_ are related to the high data quality without artifacts between 0.5 and 2 ppm. Including an MM_3co_ model reduces the 3-ppm residual by ∼30% without significant impact on the ΔAIC. All three modeling ranges (column) and three spline knot spacings (within each column), and co-edited MM models (color-coded) are presented. Bar plots represent mean values; SD is indicated by whiskers where appropriate. Minimum/maximum values for each modeling range are indicated as downwards/upwards triangles in the color corresponding to the MM_3co_ model. Minimum/maximum values for each baseline knot spacing within a specific modeling range are reported on the right side of each column. Global minimum and maximum values across all models are added as text.

The residual around the GABA+ peak (residual_ampl 3ppm_) is consistently reduced by up to 30% if a MM_3co_ model is included, in line with the reduction of structured residual in **Figure 2**. This effect is less pronounced for the dense knot spacing (0.25 ppm), indicating that a flexible baseline is to some degree capable of accounting for otherwise unmodeled MM signal. Together, these findings again support the notion that omitting an explicit MM_3co_ model does not capture the whole edited 3-ppm signal, which remains unmodeled (in the residual) or gets partially absorbed by the baseline or interpreted incorrectly as GABA signal.

The strategy with the lowest AIC is the ‘none’ model with the intermediate modeling range and sparse knot spacing, reflecting the low number of model parameters: there is no separate basis function for MM, and the low number of splines. The ΔAIC (the difference between the lowest AIC and the individual model’s AIC) consequently increases for larger modeling ranges, as more splines are included. Similarly, ΔAIC increases for denser knot spacings, and in fact, this increase is much stronger compared to the resulting reduction in both residual measures, suggesting that the increased flexibility and reduction of the residual does not justify the greater number of model parameters.

For any given knot spacing and modeling range, ΔAIC values are comparable between MM_3co_ models, with moderate increases when more parameters are estimated. Together with its low CV (9.8% compared to the minimum CV value 9.7% for the GABA_soft_ value with a narrow fit range) for GABA+, the ΔAIC for the MM09_hard_ model over the wide modeling range with sparse knot spacing (ΔAIC = 3.1) indicates a good performance of this particular model without introducing overfitting. Despite the slightly higher ΔAIC, it is beneficial to opt for the MM09_hard_ model, since the MM_0.94_ peak provides an ‘external’, non-overlapped reference anchor point for the amplitude of the expected MM_30_ peak – the MM landscape is thought to be relatively stable across healthy subjects in a narrow age range, at least in the absence of pathology ^8^. Furthermore, the MM09_hard_ model does not impose any amplitude assumptions or constraints on the target metabolite GABA.

### Separation of GABA and MM_3co_

**Figure 5** shows the distributions and CVs of the separate GABA and MM_3co_ estimates of all modeling strategies. Including a separate MM_3co_ basis function significantly decreases GABA estimates, suggesting that not doing so may lead to GABA overestimation, as MM signal is mistakenly modeled as GABA. As was seen for the composite GABA+ estimates in **Figure 3**, sparser knot spacing appears to stabilize modeling, leading to lower CVs of GABA. This becomes especially obvious for the wide modeling range, where GABA CVs exceed 50% for dense knot spacing.

**Figure 5.**
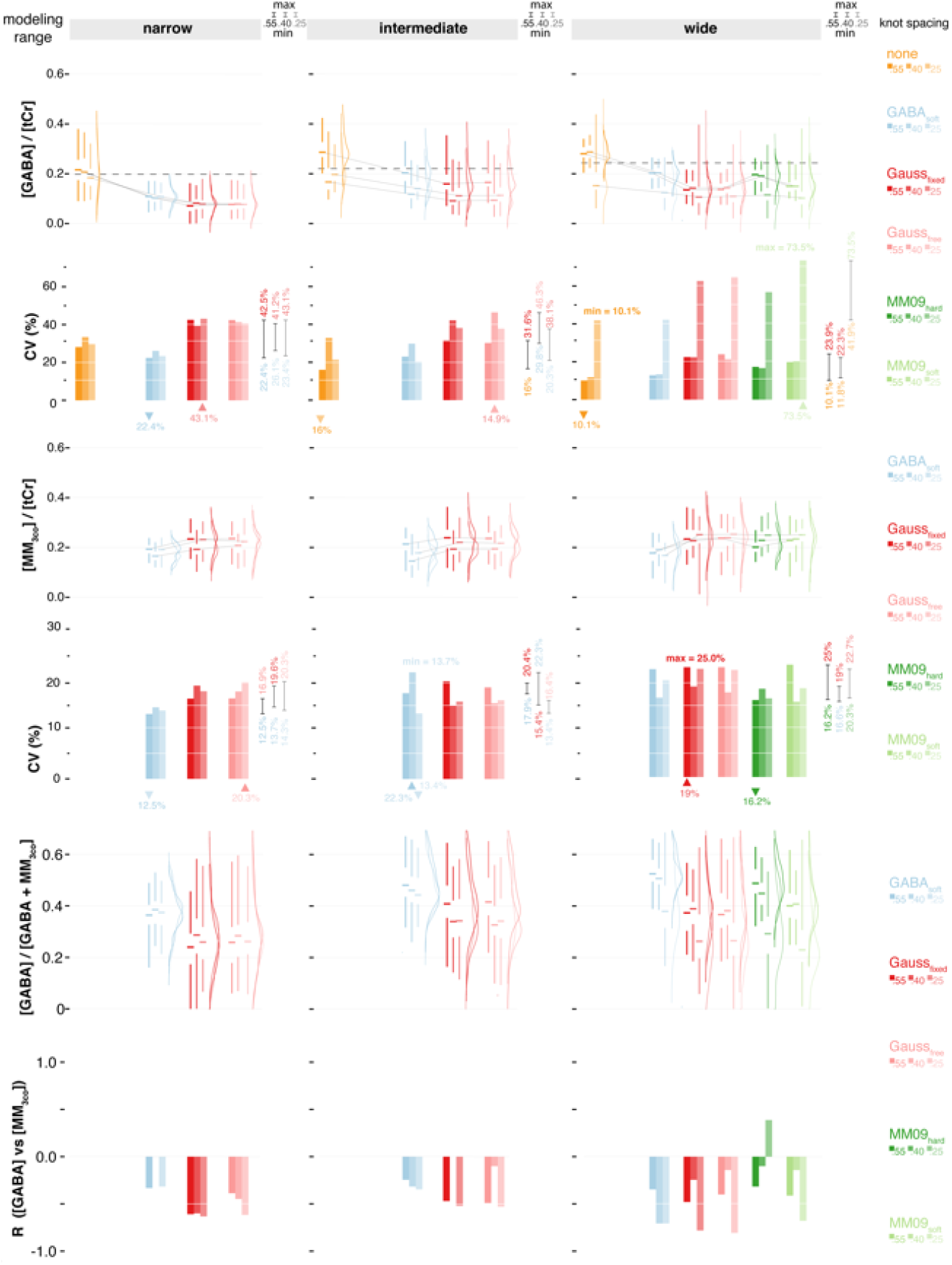
Distribution of GABA and MM_3co_ estimates, relative contribution of GABA to GABA+ and Pearson’s R between GABA and MM_3co_ for all modeling strategies. All three modeling ranges (column) and three spline knot spacings (within each column), and MM_3co_ models (color-coded) are presented. Distributions are shown as half-violins (smoothed distribution), box plots with median, interquartile range, and 25^th^/75^th^ quartile. The median lines of the box plots are connected to visualize trends within a specific baseline knot spacing. The mean estimates across the three spline knot spacings of the ‘none’ approach are indicated as a dashed line for each modeling range. Across subject CVs are summarized as bar plots. Minimum/maximum CVs for each modeling range are indicated as downwards/upwards triangles in the color corresponding to the MM_3co_ model. Minimum/maximum CVs for each baseline knot spacing within a specific modeling range are reported on the right side of each column. Global minimum and maximum CVs across all models are added as text.

MM_3co_ estimates are stable across the different knot spacings, suggesting that the different parametrizations accurately account for most of the co-edited MM signal at 3 ppm.

The GABA model, in combination with a wide modeling range and 0.55 ppm knot spacing, exhibits the lowest CV for GABA (10.4%). However, the MM09_hard_ model in combination with the same knot spacing and modeling range has only slightly higher GABA CVs (17.3%) with the corresponding MM_3co_ CVs being 16.2% (MM09_hard_). Again, despite slightly higher CV values it is beneficial to use a modeling strategy with a constraint to an ‘external’ reference peak (MM09_hard_) instead of the highly overlapped MM_3co_ peak of GABA_soft_ (CV = 12.8%), or entirely omitting MM_3co_. Additionally, the correlation between the GABA and the MM_3co_ estimates is lower for the MM09_hard_ model, potentially implying a better separation of GABA and MM_3co_. However, a separation of GABA and co-edited macromolecules remains difficult with a low-to-moderate correlation between GABA and MM_3co_ estimates for all but one modeling strategy (MM09*_hard_* for the wide fit range and 0.4 ppm baseline knot spacing). **Supplementary Material 4** and **5** reports the mean and SDs of the GABA and MM_3co_ estimates as well as the statistics, respectively.

Finally, **Figure 6** shows the impact of including HCar into the basis set with the difference in GABA and MM_3co_ estimates between the modeling strategies with and without HCar (⊗GABA and ⊗MM_3co_, respectively). Interestingly, clear differences in the systematic effects of HCar are evident between the modeling ranges:

**Figure 6.**
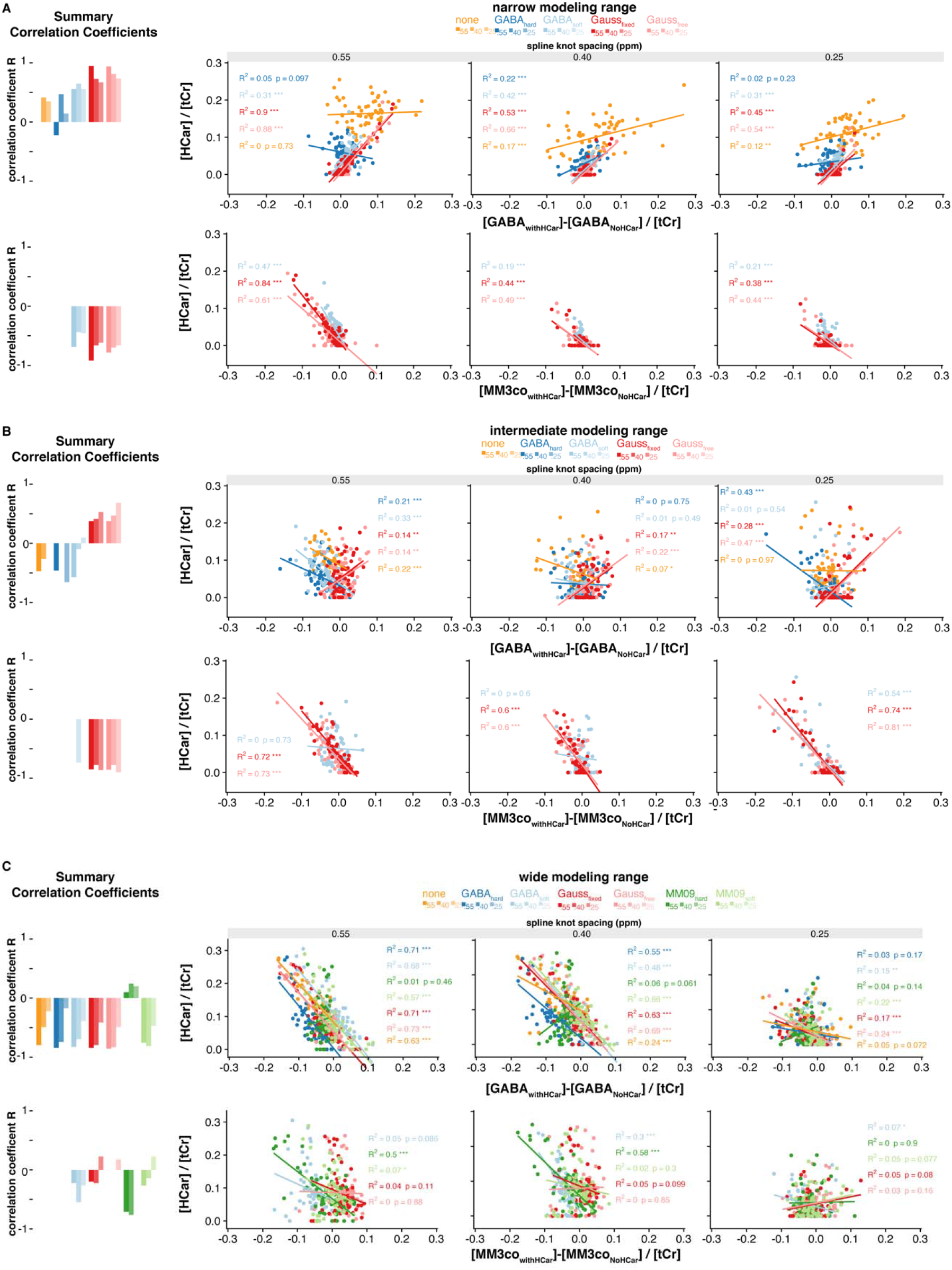
Impact of including homocarnosine in the basis set. The directionality of the correlation indicates that HCar absorbs GABA signal specifically for the intermediate and wide modeling range and absorbs MM_3co_ signal for all modeling ranges. Correlation analysis between the differences between GABA/ MM_3co_ estimates with and without HCar in the basis set and the HCar estimates. All three modeling ranges (A-C) and three spline knot spacings (within each subplot) were investigated. A summary bar plot with the correlation coefficient R is shown in the beginning of each row. Pearson’s correlation was calculated for each MM_3co_ model (color-coded). Asterisks indicate significant correlations with p < 0.05 = *, p < 0.01 = **, and p < 0.001 = ***.

For the narrow modeling range (**Figure 6 A**), HCar estimates correlate positively with ΔGABA, but the correlation is *only* substantial (R > 0.25) for strategies with a *separate* MM basis function. For precisely these strategies, HCar estimates correlate negatively with ΔMM_3co_. These observations suggest that HCar is likely to account for MM_3co_ in the narrow modeling range. In contrast, HCar and ΔGABA correlate negatively for most strategies in the intermediate and wide modeling ranges (**Figure 6 B** and **C**). The negative correlations between HCar and ΔMM_3co_ are notably weaker for these modeling ranges, indicating that HCar is more likely to substitute for GABA signal instead of MM.

This behavior can possibly be explained by the HCar signal shape for each modeling range (**Supplementary Material 6**). For the narrow modeling range, the HCar basis function offers the model an additional degree of freedom to account for deviations of the actual edited 3-ppm signal from pure GABA and the symmetric Gaussian MM3co component, as no resonances below 2.78 ppm are considered. As a result, HCar shows a high correlation with the difference in MM_3co_. For the intermediate and wide range, the HCar difference spectrum basis function more nearly resembles its GABA counterpart since other resonances are included, thereby more effectively coupling GABA and HCar estimates to each other. Perhaps unsurprisingly, HCar estimates are significantly higher for ‘none’ modeling strategy, and are substantially lower for more flexible baselines, supporting the notion that HCar rather serves as a substitute for an explicit MM signal, in particular if the baseline cannot absorb the latter (**Supplementary Material 6**). Within a given knot spacing and modeling range, HCar estimates are comparable between different MM_3co_ models, a behavior observed for GABA estimates as well. The GABA+ plus homocarnosine estimates show a slight increase compared to the GABA+ estimates without HCar (**Supplementary Material 7**). For the ‘none’ model, stronger changes occur as HCar accounts for MM signal (see also Figure 6). There was no improvement in the CVs observed when including HCar in the model. The relative contribution of HCar to GABA+ ranged between 2.2% and 19.1% for modeling strategies with an MM_3co_ basis function and between 18% and 36% for the ‘none’ model.

## Discussion

The application of linear combination modeling to edited difference spectra is neither straight-forward nor intuitive. The conceptual advantage of spectral editing arises from isolating a resolved target resonance, i.e. reducing the overlap of the target metabolite with other signals, as well as the number of signals in the spectrum in general^1^. LCM, on the other hand, benefits from maximizing the use of prior knowledge to solve the spectral modeling problem, i.e., using all available information for meaningful constraint, including from overlapping signals. The specific case of GABA-edited MRS at 3T poses unique and unresolved challenges. Firstly, a compromise must be drawn between maximizing the prior knowledge by increasing the modeling range and reducing the impact of co-edited and unwanted signals. Secondly, an appropriate parametrization of poorly characterized co-edited signals must be found, and possible interactions with the target metabolite GABA must be evaluated. Thirdly, effects of baseline modeling must be studied, again a consequence of the macromolecular background signal in the GABA-edited difference spectrum not being determined to this date. In this study, a total of 102 linear combination modeling strategies were compared for GABA-edited difference spectra, each with different modeling ranges, parametrizations of co-edited signals, and baseline model flexibility. The key findings are:

- Including a dedicated basis function for co-edited MM improves fit residuals, reduces CVs of GABA and GABA+ estimates, and avoids overestimation of GABA.
- Reducing the modeling range does not substantially stabilize or destabilize modeling, while removing potentially valuable information (MM_0.93_ and 2-ppm NAA peak) from the optimization.
- Sparser baseline spline knot spacing leads, on average, to the lowest CV across all modeling ranges.

There is surprisingly little systematic investigation into linear combination modeling of GABA-edited difference spectra. To the best of our knowledge, there is only one conference abstract studying MM parametrization in GABA-edited MRS with the LCModel software^27^. The results from this preliminary investigation indicate that including a specific MM basis function significantly reduces GABA estimates, as was also observed in an earlier study ^28^ and which is substantiated by our findings.

Although the substantial contribution of broad MM signals to the 3-ppm peak in the GABA-edited spectrum is widely known^1,35^, it is rarely explicitly addressed in linear combination modeling. Instead, it is assumed that either an incomplete model (without explicit MM term) will still provide an accurate GABA estimate, or that baseline modeling will account for the MM signal. The current results provide evidence that including an appropriately parametrized MM model is a preferrable and easily implemented strategy, reducing the residual over the 3-ppm signal range by up to 30%, with similar or lower CVs for GABA+. In contrast, not including an MM model likely causes systematic overestimation of GABA, as the least-squares optimization attempts to minimize the model-data difference with an inadequate set of basis functions (only GABA), particularly when a rigid baseline is chosen. Including MM_3co_ is a justified and reasonable measure without overfitting (reflected by AIC), and stable mean estimates and CVs of MM_3co_ suggest an adequately parametrized model. In addition, it is notable that including MM_3co_ is increasingly beneficial for the narrow fit range as it leads to a significant reduction in the SD of the GABA+ estimates. This SD reduction is overserved for three models (Gauss_fixed_, Gauss_free_, MM09_hard_) for the wide fit range with 0.55 ppm baseline knot spacing and not overserved for the intermediate fit range.

The different MM models in this study were based on certain assumptions, including the relative contribution of MM_3co_ to the 3-ppm GABA peak to be around 50%^1,6,8,19^. Levels of MM_0.93_ have been found to be stable across the whole brain^36^ and are thought to be stable across healthy subjects. Under these assumptions, the MM09_hard_ model with a rigid amplitude coupling between MM_3co_ and the non-overlapped MM_0.93_ peak is a suitable strategy, supported by favorable CVs and ΔAIC. Further studies need to be performed to investigate the distribution and correlation between MM_0.93_ and MM_3co_ in the brain. For the MM09soft modeling strategy, we have found the following MM_3co_/MM_0.93_ ratios: 0.97 ± 0.29 (0.55 ppm baseline knot spacing), 0.88 ± 0.18 (0.4 ppm baseline knot spacing), and 0.95 ± 0.20 (0.25 ppm baseline knot spacing) compared to 0.66 for the composite MM09_hard_ model. This indicates higher ratios then expected in the initial model parameters. However, true values can only be inferred from a large number of measured macromolecular background spectra.

Compared to the Gauss_fixed_ model, the Gauss_free_ approach has one additional parameter to change the FWHM of the MM_3co_ basis function. However, it can be assumed that difference in the linewidth would mostly be accounted for by the Lorentzian linebroadening term. Therefore, only minor differences in the fit results are expected, especially considering the high data quality in this study. This was indeed the case in this study. As an example, for the wide fit range and 0.55 ppm baseline knot spacing, the FWHM of the MM_3co_ basis function was 14 Hz for the Gauss_fixed_ model and 14.01 ± 0.10 Hz for the Gauss_free_ model.

Unedited MRSI data measured at 7T indicates significant differences between white and gray matter for several macromolecules in the healthy brain^36^. Changes in the MM concentrations during disease may also affect the relative contribution to the 3-ppm peak, and therefore render models with prior amplitude assumptions inaccurate. If there is reason to expect strong fluctuations of MM_3co_, a modeling strategy with fewer assumptions about amplitude ratios between the metabolite of interest GABA or the MM_0.93_ signal and the MM_3co_ signal is preferable to the MM09_hard_ strategy. Here, the Gauss_free_ and Gauss_fixed_ strategies could be used to account for changes in the MM_3co_ contribution more freely, as their mean estimates of GABA and GABA+ were in good agreement with the more constrained approaches, although they led to increased CVs and ΔAICs. In addition, the less-constrained models might be more appropriate for investigating changes in MM_3co_ due to age^37^ or disease, or for exploring frequency-drift-related effects on the co-edited MM signal^1,8,38^. Another potential way to model the co-edited MM signal is to include lysine in the simulated basis set, as it has been identified as the potential source of the signal^6^, although this approach would require appropriate broadening and incorporation of chemical shift and coupling values from protein databases^39^.

Overall, results did not differ drastically between modeling ranges, although it is noteworthy that the effects of baseline flexibility were less pronounced for the narrow modeling range, likely because the complex interaction of the overlapping 2.25 ppm GABA and Glx signals with the underlying baselines is omitted. Furthermore, there was no evidence that the intermediate modeling range, which is proposed in the LCModel manual^18^ to avoid frequently occurring co-edited lipid signals, improved quantification substantially compared to both other modeling ranges, although it should be mentioned that this particular dataset did not suffer from severe lipid contamination. Taken together, the choice of modeling range does not impact quantitative results as substantially as the inclusion of an MM model.

Baseline models are included in most LCM algorithms to account for signals not otherwise modeled, e.g., residual water tails or unparametrized macromolecules and lipids. Compared to conventional short-TE spectra, water and non-co-edited MMs are removed upon subtraction in the GABA-edited spectrum, which is therefore frequently modeled with a stiffer baseline^4,18^. Our results show that sparser knot spacing (0.55 ppm) leads to lower CVs in metabolite estimates. A more flexible baseline (0.25 ppm) improves local and global residuals, but not enough to justify the additional model parameters (as per the AICs). More importantly, an overly flexible baseline may absorb edited signal, although it appeared that it did not do so excessively even for the 0.25-ppm strategies. The exception was the ‘none’ model, where the baseline was the only available part of the model to take up signal, underlining the inadequacy of the default LCModel approach. Taken together, a relatively rigid baseline with a parametrized MM basis function is preferable for LCM of GABA-edited spectra. A caveat to this recommendation is the observation of structural baseline fluctuations underneath the 2.25 ppm signals from GABA, Glx, GSH, NAA and NAAG, particularly for the 0.25 ppm knot spacing and a relatively broad increase in the baseline between 2.7 and 3.3 ppm. These were observed previously^27^, and are likely signals from unparametrized MMs directly and indirectly affected by the editing pulse. Rigid baselines may force a wrong metabolite model in that region and interfere with accurate estimation of GABA and Glx. In fact, the structural Glx residual at 3.75 ppm suggests a systematic misestimation of the Glx phase, likely driven by the 2.25 ppm signals. While beyond the scope of this investigation, it is conceivable that more informed parametrization (or, ideally, direct measurement) of this unexplored MM background may benefit the modeling of the entire difference spectrum. Alternatively, hitherto unexplored approaches with variable baseline knot spacing may be worth investigating.

The HCar molecule has a GABA moiety with similar chemical shifts and is therefore co-edited. Evidence regarding in-vivo HCar levels in the human brain is inconclusive – early work determined HCar levels to be 0.5 mM^5^ (compared to ∼1 mM for GABA), while a recent hybrid up-field/downfield inversion-recovery method determined the HCar/GABA ratio as 17%^40^. Therefore, we tested the impact of adding HCar to the basis set without additional constraints. Including HCar systematically affected GABA and MM_3co_ estimates, in a way that strongly depended on the choice of modeling range. HCar estimates themselves ranged from 2.2% to 19.1% of the GABA+ signal, depending strongly on the degree of baseline flexibility. The results suggest that the overlap between the three model terms (HCar, GABA, MM_3co_) is too substantial for reliable three-way separation, particularly in the presence of a highly flexible baseline. A minor increase in “GABA+ plus HCar” estimates compared to GABA+ estimates was observed and the inclusion of HCar did not substantially improve the CVs. Additionally, the disagreement between the model and the data at 2.9 ppm indicates that a simple unconstrained addition of HCar to the modeling is not justified.

Symmetric GABA-editing (edit-ON frequency at 1.9 ppm and edit-OFF frequency at 1.5 ppm) is commonly used eliminate the MM_3co_ contamination of the 3-ppm GABA+ signal. In practice, B_0_ instabilities lead to residual MM_3co_ components with variable polarity ^11^. The Gauss_free_ and Gauss_fixed_ MM_3co_ models could potentially be used to account for those variable MM_3co_ contributions in those spectra. However, modeling of those spectra with the current strategies that do have a non-negative model component as constraint would be challenging. Those modeling strategies could potentially be adapted by using the B_0_ history during the experiment ^38^ to predict the polarity and relative amplitude of the MM_3co_ signal, and include those as a soft constraint relative to the MM_09_ signal (MM09_hard_ or MM09_soft_) or the GABA signal (GABA_hard_ or GABA_soft_).

### Limitations

A limitation of this study is the high spectral quality (SNR, linewidth, no apparent subtraction artefacts, or lipid contaminations) of the dataset analyzed. We did not investigate model parametrizations of movement or drift, which may introduce systematic changes to the co-edited MM signal. While our results suggest that using the wide modeling range with a rigid baseline is beneficial, strong co-edited lipid signals are likely to not be modeled appropriately, and the intermediate modeling range may be more suitable. Further studies of the possible impact of changes in spectral quality need to be performed to validate the modeling strategies under suboptimal conditions.

Another limitation is that there is no ‘gold standard’ of metabolite level estimation in GABA-edited MRS to validate the results against. The performance of different algorithms or in this study modeling strategy is often judged by the level of variance ^26^. A lower variance does, of course, not necessarily reflect greater modeling accuracy, but under the assumption that the homogeneous study population and data acquisition contribute comparably little biological and instrumental variance, CVs will predominantly reflect variance introduced by the modeling approach. Recently, the field is witnessing increasing efforts to generate simulated spectra with known ground truth as a gold standard, although these approaches can only be successful to the extent that those spectra are truly representative of in-vivo data ^41–43^. Further, such gold standard studies with a known ground truth could be used to validate whether a correct separation of GABA and MM_3co_ is achievable by advanced LCM. This study indicates a low-to-moderate correlation between the GABA and MM_3co_ estimates, suggesting that the two components are not reliably separated. However, some of the modeling strategies appeared to have a lower association between both estimates and could possibly be validated further on a synthetic dataset with known GABA and MM_3co_ concentrations.

AIC as a measure of the goodness of fit can be used for linear and non-linear approaches if the log-likelihood is obtained similarly. However, there are two potential limitations for the application of AIC in this study. First, for linear-combination modeling of MRS data, as implemented in Osprey, a non-linear optimization is followed by a linear optimization during each iteration. Parameters are treated equally in the calculation of the AIC regardless of whether they are non-linear (e.g., a phase parameter) or linear (an amplitude parameter). Second, the AIC penalizes complex models, but does not measure effects of soft constraints and is likely to prefer models without a soft constraint as those should have a reduced likelihood ^44^. Here, we introduced a rather arbitrary correction term of 0.5 per soft constraint for those models to reduce this effect. Therefore, the resulting ΔAIC values in this study should be interpreted with care and considered as only one among several metrics to evaluate model performance.

## Conclusion

This study proposed and compared different modeling strategies for LCM of GABA+-edited difference spectra from a multi-site MEGA-PRESS dataset. Introducing a parametrized model for co-edited macromolecules reduces fit residuals, while maintaining low coefficients of variation of GABA+ estimates. A rigid baseline was found to be beneficial, while using a narrower modeling range did not significantly improve the modeling. The overall modeling results suggest that GABA-edited data are reliably modeled with an adequately parametrized MM_3co_ model, constrained by the non-overlapped 0.93-ppm MM resonance, in combination with a full modeling range (between 0.5 and 4 ppm) and sparse knot spacing (0.55 ppm). Incorporating homocarnosine into the modeling did not significantly improve the GABA+ estimates and did not allow for a stable separation of GABA and HCar.

## Supplementary Material

### Table of Contents

1. List of included subjects
2. Macro-molecule function definitions
3. Mean and SD spectra as well as individual spectra
4. Statistics - GABA estimates
5. Statistics - MM_3co_ estimates
6. Model overview basis set with homocarnosine
7. Distribution of GABA+ and homocarnosine

**Supplementary Material 1.** – List of included subjects. All datasets are available at https://www.nitrc.org/projects/biggaba/

| site | subjects | $\Sigma$ |
| --- | --- | --- |
| P01 | S01,S03,S04,S05,S08 | 5 |
| P03 | S02,S03,S04,S07,S08,S09,S10,S11,S12 | 9 |
| P05 | S01,S02,S03,S05,S06,S07,S08 | 7 |
| P06 | S01,S02,S03,S04,S05,S06,S07,S08,S09 | 9 |
| P07 | S02,S03,S04,S09,S10,S11,S12 | 7 |
| P08 | S01,S02,S03,S04,S05,S06,S07,S08,S09,S10,S11,S12 | 12 |
| P09 | S01,S02,S03,S04,S05,S06,S07,S08,S09,S10,S11,S12 | 12 |
| $\Sigma = 7$ | | $\Sigma = 61$ |

**Supplementary Material 2.** Properties of the Gaussian functions of the broad macromolecule and lipid resonances included in the basis sets, taken from section 11.7 of the LCModel manual. The amplitude values are scaled relative to the CH_3_ singlet of creatine with amplitude 3.

| Name | Frequencies [ppm] | FWHM [ppm] | Amplitude |
| --- | --- | --- | --- |
| *edit-OFF spectrum basis set* |  |  |  |
| MM<sub>0.94</sub> | 0.915 | 0.14 | 3.00 |
| MM<sub>1.22</sub> | 1.22 | 0.15 | 2.00 |
| MM<sub>1.43</sub> | 1.43 | 0.17 | 2.00 |
| MM<sub>1.70</sub> | 1.67 | 0.15 | 0.20 |
| MM<sub>2.05</sub> | 2.08 | 0.15 | 1.33 |
|  | 2.25 | 0.20 | 0.33 |
|  | 1.95 | 0.15 | 0.33 |
|  | 3.00 | 0.20 | 0.40 |
| Lip09 | 0.89 | 0.14 | 3.00 |
| Lip13 | 1.28 | 0.15 | 2.00 |
|  | 1.28 | 0.089 | 2.00 |
| Lip20 | 2.04 | 0.15 | 1.33 |
|  | 2.25 | 0.15 | 0.67 |
|  | 2.80 | 0.20 | 0.87 |
| Difference spectrum basis set |  |  |  |
| MM<sub>0.94</sub> | 0.915 | 0.14 | 3 |
| MM<sub>3co</sub> | 3 | **14 Hz** | 2 |

**Supplementary Material 3.**
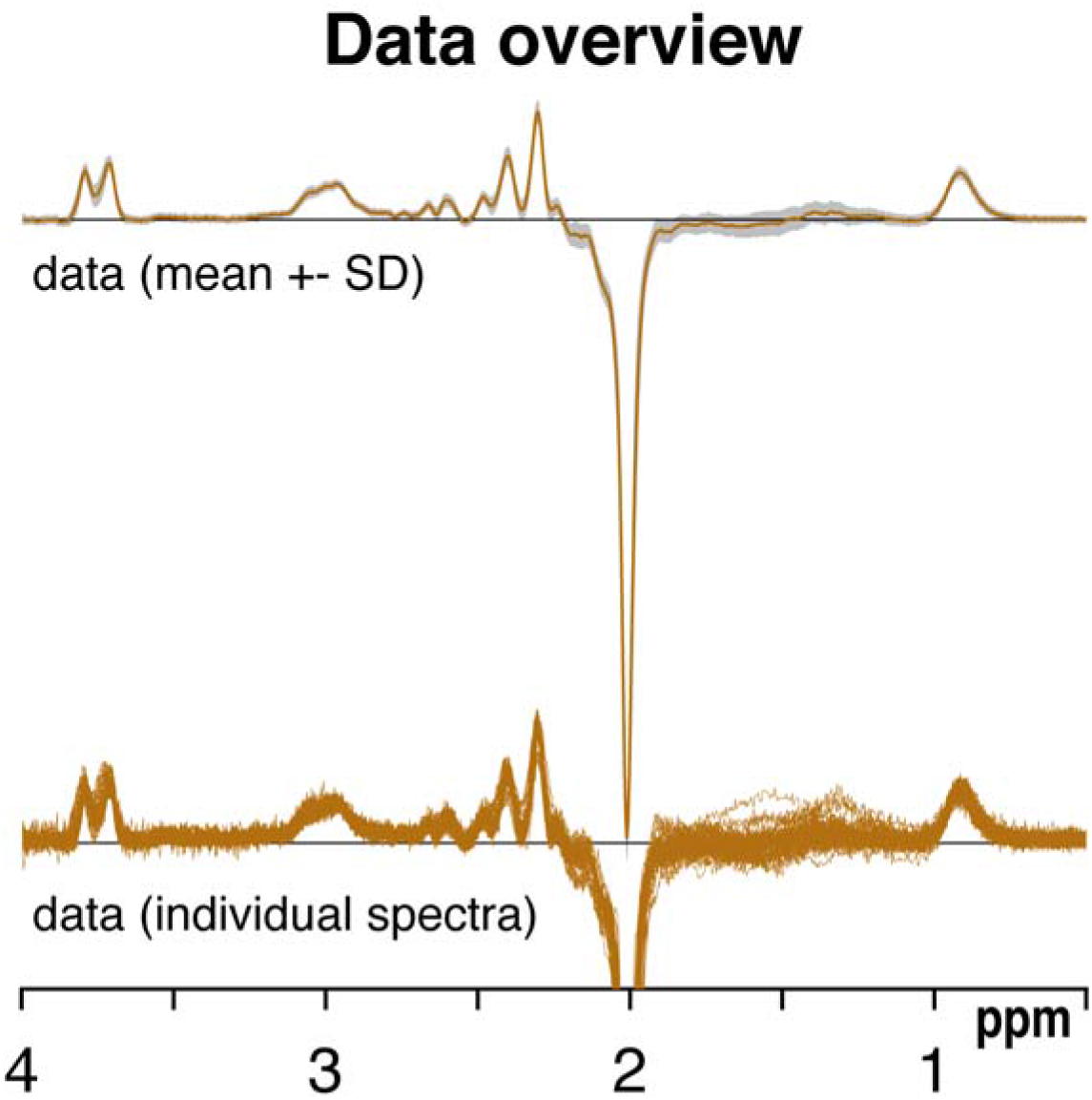
Overview of the processed data including the mean ± SD and individual data.

**Supplementary Material 4.** GABA mean and SDs for all modeling strategies (ratios to tCr). Significant differences (p < .05) between the corresponding model and the ‘none’ model (gray shade) are indicated in bold.

| modeling range |  | narrow |  |  | intermediate |  |  | wide |  |  |
| --- | --- | --- | --- | --- | --- | --- | --- | --- | --- | --- |
| knot spacing (ppm) |  | 0.55 | 0.4 | 0.25 | 0.55 | 0.4 | 0.25 | 0.55 | 0.4 | 0.25 |
| none | [GABA] | 0.220<br>.062 | 0.213<br>.071 | 0.193<br>.057 | 0.297<br>.048 | 0.186<br>.061 | 0.204<br>.044 | 0.284<br>.029 | 0.288<br>.034 | 0.158<br>.066 |
| GABA<sub>hard</sub> | [GABA] | **0.164**<br>**.021** | **0.146**<br>**.02** | **0.156**<br>**.023** | **0.207**<br>**.024** | **0.148**<br>**.031** | **0.165**<br>**.021** | **0.194**<br>**.021** | **0.207**<br>**.023** | **0.150**<br>**.034** |
| GABA<sub>soft</sub> | [GABA] | **0.108**<br>**.024** | **0.102**<br>**.027** | **0.108**<br>**.025** | **0.207**<br>**.048** | **0.130**<br>**.039** | **0.148**<br>**.03** | **0.203**<br>**.026** | **0.204**<br>**.027** | **0.115**<br>**.049** |
| Gauss<sub>fixed</sub> | [GABA] | **0.074**<br>**.031** | **0.080**<br>**.031** | **0.081**<br>**.035** | **0.174**<br>**.055** | **0.106**<br>**.045** | **0.118**<br>**.045** | **0.136**<br>**.031** | **0.145**<br>**.032** | **0.108**<br>**.068** |
| Gauss<sub>free</sub> | [GABA] | **0.079**<br>**.034** | **0.078**<br>**.032** | **0.081**<br>**.033** | **0.180**<br>**.054** | **0.107**<br>**.050** | **0.119**<br>**.045** | **0.137**<br>**.033** | **0.145**<br>**.031** | **0.107**<br>**.069** |
| MM09<sub>hard</sub> | [GABA] | - | - | - | - | - | - | **0.195**<br>**.034** | **0.189**<br>**.032** | **0.106**<br>**.060** |
| MM09<sub>soft</sub> | [GABA] | - | - | - | - | - | - | **0.152**<br>**.03** | **0.154**<br>**.031** | **0.098**<br>**.072** |

**Supplementary Material 5.**
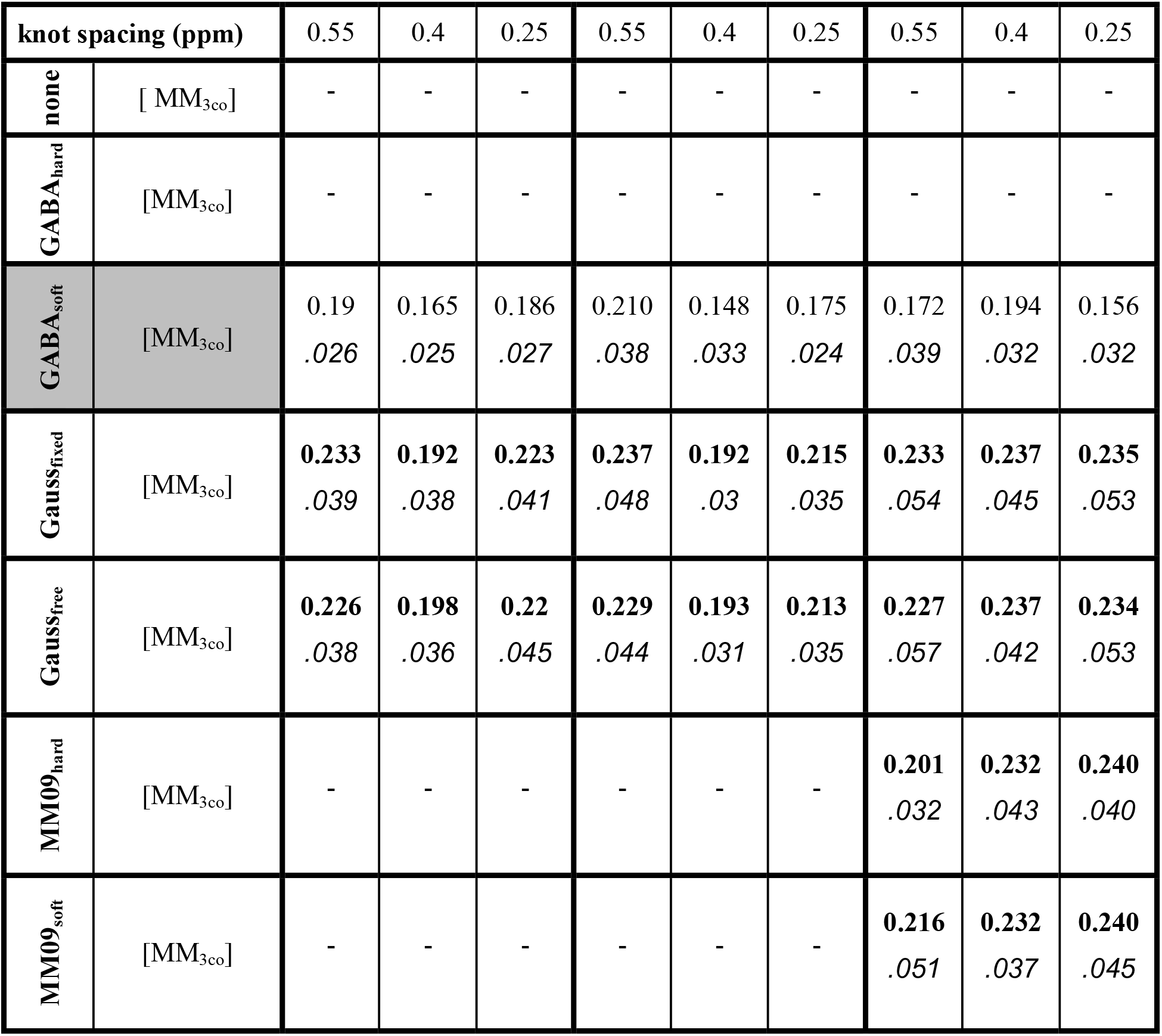
MM_3co_ mean and SDs for all modeling strategies (ratios to tCr). Significant differences (p < .05) between the corresponding model and GABA_soft_ model (gray shade) are indicated in bold.

| knot spacing (ppm) |  | 0.55 | 0.4 | 0.25 | 0.55 | 0.4 | 0.25 | 0.55 | 0.4 | 0.25 |
| --- | --- | --- | --- | --- | --- | --- | --- | --- | --- | --- |
| none | [MM<sub>3co</sub>] | - | - | - | - | - | - | - | - | - |
| GABA<sub>hard</sub> | [MM<sub>3co</sub>] | - | - | - | - | - | - | - | - | - |
| GABA<sub>soft</sub> | [MM<sub>3co</sub>] | 0.19<br>.026 | 0.165<br>.025 | 0.186<br>.027 | 0.210<br>.038 | 0.148<br>.033 | 0.175<br>.024 | 0.172<br>.039 | 0.194<br>.032 | 0.156<br>.032 |
| Gauss<sub>fixed</sub> | [MM<sub>3co</sub>] | 0.233<br>.039 | 0.192<br>.038 | 0.223<br>.041 | 0.237<br>.048 | 0.192<br>.03 | 0.215<br>.035 | 0.233<br>.054 | 0.237<br>.045 | 0.235<br>.053 |
| Gauss<sub>free</sub> | [MM<sub>3co</sub>] | 0.226<br>.038 | 0.198<br>.036 | 0.22<br>.045 | 0.229<br>.044 | 0.193<br>.031 | 0.213<br>.035 | 0.227<br>.057 | 0.237<br>.042 | 0.234<br>.053 |
| MM09<sub>hard</sub> | [MM<sub>3co</sub>] | - | - | - | - | - | - | 0.201<br>.032 | 0.232<br>.043 | 0.240<br>.040 |
| MM09<sub>soft</sub> | [MM<sub>3co</sub>] | - | - | - | - | - | - | 0.216<br>.051 | 0.232<br>.037 | 0.240<br>.045 |

**Supplementary Material 6.**
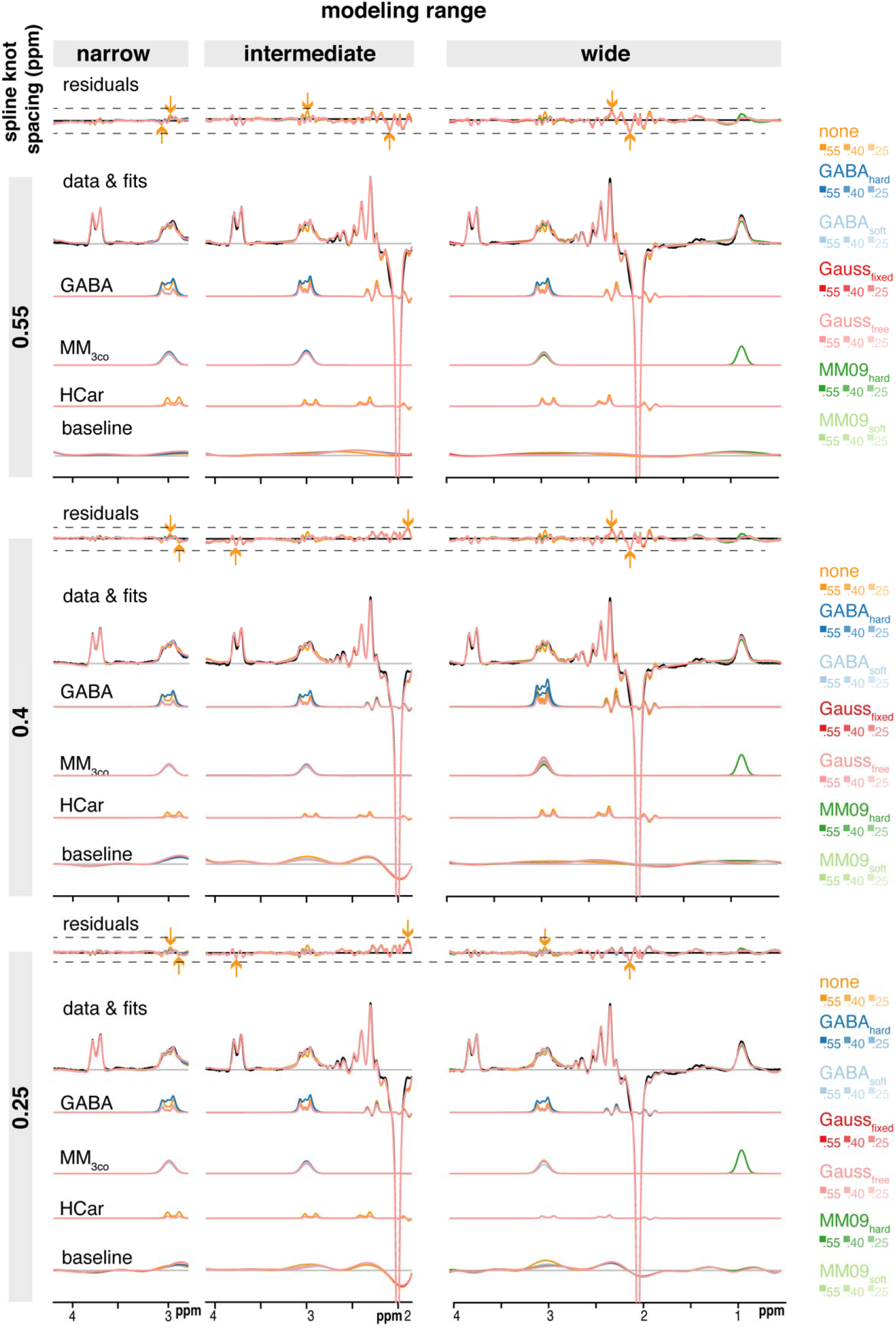
Mean modeling results and homocarnosine estimates for all modeling strategies with homocarnosine. A substantial structured residual is visible at 3 ppm if for all modeling strategies and for the narrow and intermediate modeling range the homocarnosine concentrations are significantly lower compared to omitting the co-edited MM, especially for knot spacings <= 0.4 ppm. All three modeling ranges (columns), three spline knot spacings (rows), and MM_3co_ model (color-coded) are presented with mean residuals and fits, as well as the GABA, MM_3co_, homocarnosine (HCar) and spline baseline models. The mean data is included in black. The arrows indicate the range of values for a specific modeling range and spline knot spacing with the color corresponding to the MM_3co_ model with minimum/maximum value.

**Supplementary Material 7.**
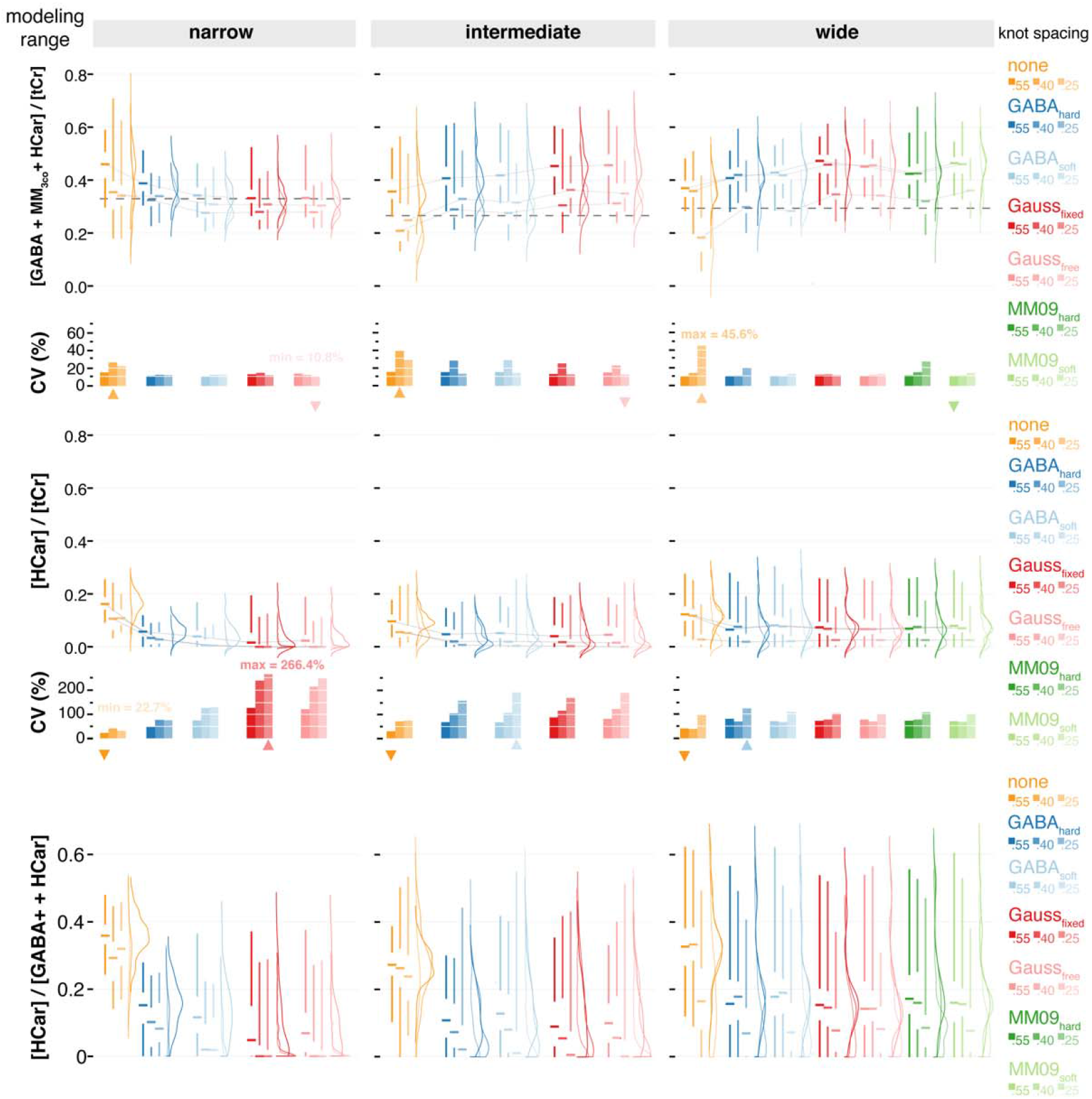
Distribution of GABA+ plus HCar and HCar estimates and the relative contribution of HCar to GABA+ plus HCar for all modeling strategies. The mean estimates of GABA+ plus HCar across the three spline knot spacings of the ‘none’ approach are indicated as a dashed line for each modeling range. All three modeling ranges (column) and three spline knot spacings (within each column), and MM_3co_ models (color-coded) are presented. Distributions are shown as half-violins (smoothed distribution), box plots with median, interquartile range, and 25^th^/75^th^ quartile. The median lines of the box plots are connected to visualize trends within a specific baseline knot spacing. CVs are summarized as bar plots. Minimum/maximum CVs for each spline knot spacing are indicated as downwards/upwards triangles in the color corresponding to the MM_3co_ model. Global minimum and maximum CVs across all models are added as text.

## Acknowledgement

This work is supported by NIH grants P41 EB031771, R01 EB016089, R01 EB023963, R01 EB028259, R21 AG060245, and K99/R00 AG062230.

## CRediT authorship contribution statement

**Helge J. Zöllner**: Software, Formal Analysis, Investigation, Writing – Original Draft, Writing – Review & Editing, Visualization. **Sofie Tapper**: Investigation, Writing – Review & Editing. **Steve C. N. Hui**: Investigation, Writing – Review & Editing. **Richard A. E. Edden**: Conceptualization, Formal Analysis, Writing – Review & Editing, Supervision, Project administration, Funding acquisition. **Peter B. Barker**: Writing – Review & Editing, Supervision, Funding acquisition. **Georg Oeltzschner**: Conceptualization, Methodology, Software, Investigation, Formal Analysis, Writing – Review & Editing, Supervision, Funding acquisition.

